# Cytokinin-induced immunity enhances rice blast resistance

**DOI:** 10.1101/2024.05.10.593599

**Authors:** Emilie Chanclud, Anna Kisiala, R.J. Neil Emery, Grégory Mouille, Stéphanie Boutet, Corinne Romiti-Michel, Jean-Benoit Morel

## Abstract

Plant cytokinins (CKs) affect the outcome of plant-pathogen interactions. However, despite many examples their role remains ambiguous. In rice, CKs act synergistically with salicylic acid (SA) to induce defense genes *in vitro* but no effect on resistance against the blast fungus, *Magnaporthe oryzae,* was further observed *in planta*. Here, we demonstrate that exogenous CKs treatment triggers rice blast resistance in a molecule-, dose- and time-dependent manner by affecting defense- and CK-responsive genes. Similar enhanced resistance and gene expression patterns were confirmed in rice insertion mutant lines impaired for a gene that encodes a putative CK inactivation enzyme, supporting that endogenous CKs affect rice immunity. Together, our work brings insights on the CK-induced resistance of rice to *M. oryzae*.

## Introduction

Plant hormones are key players to fine-tune plant physiology and developmental adaptations to abiotic and biotic environments ^1–4^. Hormonal homeostasis allows a wide range of responses and its precision requires multiple cross-talks between their metabolic and response pathways ^2,5–7^. Overall, this makes it difficult to unravel their individual impacts by exogenous treatments or by genetic mutations ^8–10^. Cytokinins (CKs) are adenine derivatives compounds that were first described for their effects on plant physiological processes (i.e. cellular division and differentiation ^11,12^, photosynthesis and senescence ^13,14^, nutrient source/sink relationships ^15,16^) and for being a key communication factor in the establishment of the bacterial symbiosis ^17,18^. Today, along with their wide roles in mutualisms ^19,20^, CKs are also well implicated to impact outcomes of plant-pathogen interactions ^1,21–29^. Their variable effects on plant resistance often depend on the host and on the pathogen’s trophic behavior ^5^. Several studies showed that CKs act synergistically with salicylic acid (SA) to induce plant defenses and that the SA pathway is required for the CK-triggered immunity ^9,22,25,29^. Additionally, CKs are acknowledged to affect microbial development and physiology ^30,31^. However, even if their role in the virulence of the tumor-inducing pathogens is well understood it remains mostly elusive in the case of the non-tumor inducing ones ^31–36^.

To unravel CKs involvement in such interactions, the rice blast disease caused by *Magnaporthe oryzae* was proposed as a good model system ^9,24^. Indeed, previous *in vitro* studies demonstrated that in rice, CKs and SA act synergistically to induce defense marker genes expression but without any effects on resistance *in planta* ^9^. In the present study, to further explore how CKs affect rice immunity we investigated the impact of exogenous CK-applications on blast resistance. To do so, we quantified symptoms, fungal penetration events and measured defense- and CK-marker genes expression upon infection, after the plants were sprayed with CKs-mimicking compounds (kinetin - KIN - and benzyladenine - BA). We varied the concentrations (from 10µM to 500µM) and the timing of application before inoculation (24h, 48h, 72h) with different virulent isolates (GY11 moderately virulent and FR13 virulent, on Nipponbare rice). This enabled us to define conditions that brought out a reproducible CK-induced resistance phenotype. By reverse genetics, we subsequently confirmed that endogenous CKs similarly induce immunity of rice CK mutants. Together, our results support that under specific conditions, CK-induced immunity could contribute to rice blast resistance.

## Results

### Exogenous CKs trigger rice blast resistance in a compound-, time-, and dose-dependent manner

Plants treated with KIN (50µM) 48h before inoculation (hbi) were more resistant to rice blast (Fig. 1; Suppl. Fig. S1). We observed a reduction of the number of sporulating lesions by about 30% compared to the untreated plants (Fig. 1AB; Suppl. Fig. S1AB). The microscopic observations confirmed a reduced pathogen growth inside the host tissues with a lower number of spores forming the primary hyphae (Fig. 1C; Suppl. Fig. S1C). However, no difference in resistance was observed among plants treated earlier than 48 hbi (24hbi) with 50 µM KIN or BA (Suppl. Fig.S2AB).

**Figure 1.**
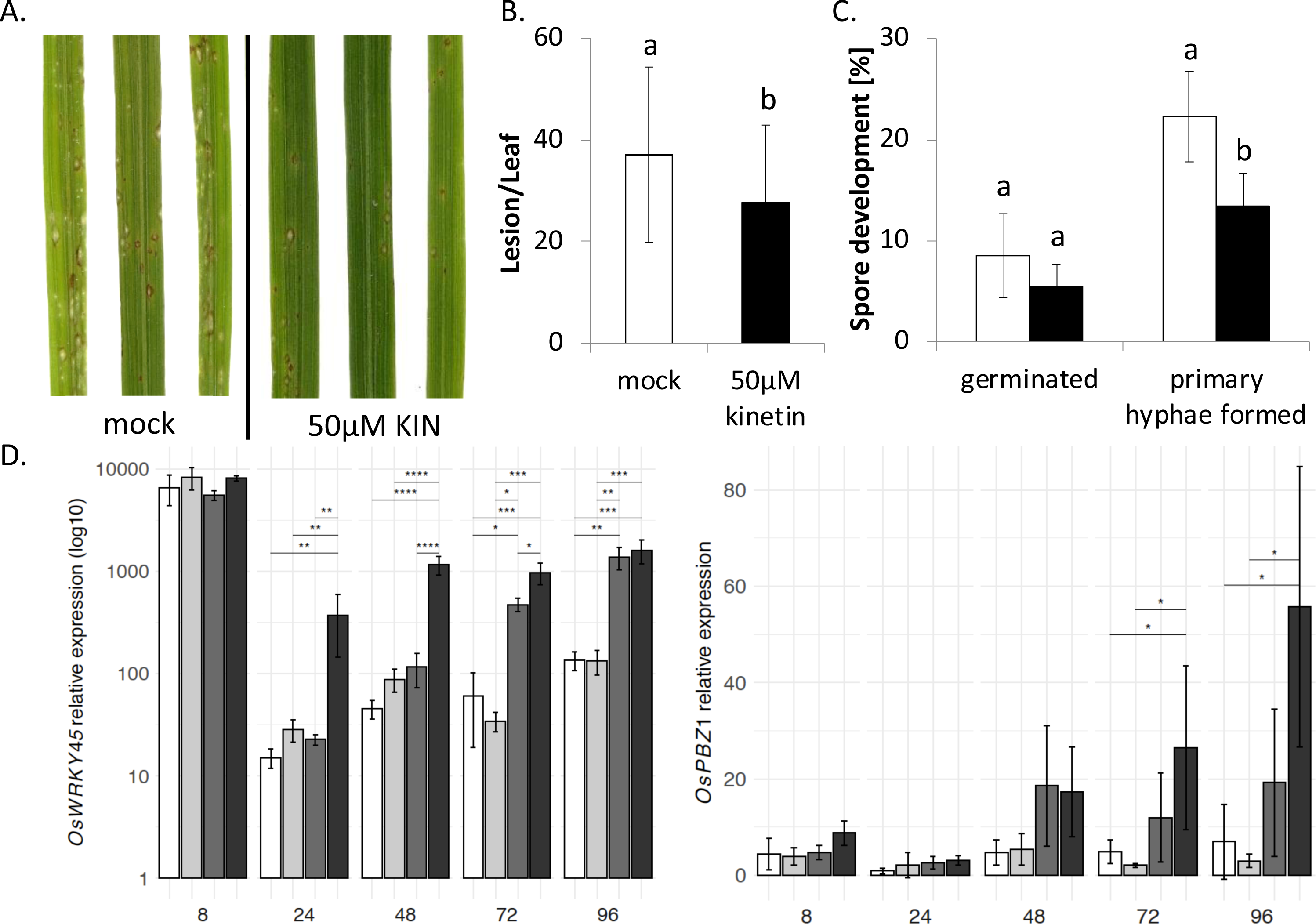
Kinetin enhances rice blast resistance. (A) Symptoms caused by the *M. oryzae* strain GY11 on Nipponbare rice plants treated with 50µM of kinetin (KIN) 48h before infection (hbi) and on the controls (mock) observed 6 days after inoculation. (B) Symptom quantification. The values represent the mean and SD from six biological replicates each composed of 10 plants (t-test, p-value = 0.02). (C) Microscopic observations of spore germination and penetration 72h post inoculation in plants pretreated with 50µM of KIN 48hbi (black bars) and in the controls (mock; white bars). The data presented is the mean and SD of three biological replicates (>50 infection sites/replicate, t-test, p-value = 0.02). (D) Relative expression of defense-marker genes, *Os WR K Y 45* and *PBZ 1*, in mock-treated plants sprayed with 0.5% gelatin (white bars), in KIN-treated plants sprayed with 0.5% gelatin (light grey bars), in mock-treated infected plants (dark grey bars) and in KIN-treated infected plants (black bars) at 8, 24, 48, 72 and 96h after inoculation (hai). The transcriptional regulation was evaluated by quantitative RT-PCR using the *Actin* gene for normalization. The values presented are the means and SD calculated from four independent replicates composed of 6 plants. Asterisks indicate significance in a one-way analysis of variance with a Tukey post hoc test, *p* < 0.04. For all the experiments, KIN 50mM in 50% EtOH diluted x1000 (KIN 50µM final) or the equivalent 50% EtOH control mock solution were applied to plants 48hbi. Inoculated plants were sprayed with spore suspension (in gelatin 0.5%) and uninfected controls with gelatin (0.5%) only.

Using the application time point that gave the most reproducible results (48hbi), we further investigated the dose-effect of KIN (10-500µM) on Nipponbare resistance to the GY11 isolate (Suppl. Fig. S3). All tested concentrations of KIN above 10µM enhanced or tended to enhance resistance, whereas 50µM was the lowest concentration that triggered a reproducible and significant phenotype.

### Exogenous KIN enhances SA-related marker genes express ion upon infection

To test the hypothesis that KIN (50µM, 48hbi) synergistically induces defense genes with the SA-pathway triggered by the infection, we measured the expression of the SA-responsive defense-marker genes, *Os WR K Y 45* and *Os PBZ 1,* and of SA-independent defense markers (including *SP L 7*, *POX 223*, *PR 5*; Fig. 1D and Suppl. Fig. S1D, Suppl. Fig. S4). The expression of most of these genes was significantly higher during infection in KIN-treated plants compared to the controls (Fig, 1D, Suppl. Fig. S1D, Suppl. Fig. S4).

### Exogenous KIN and BA trigger different CK-marker genes res ponses

Physiological CKs responses rely in part (i) on the pool of active forms, which depends on their degradation/inactivation and (ii) on the downstream transcription responses involving CK-response regulators (RR). Here, we hypothesized that the effectiveness of the exogenous CK treatments observed for different compounds, doses and timing coincides with the varying induction of the downstream CK-response pathways. To test this hypothesis, we measured the expression of some CK-related marker genes (such as *C ytokinin Oxidase* (CKX) and CK *RR* genes), in plants infected and pre-treated with 50µM of KIN or BA at different hbi (Suppl. Fig. S2C). Matching with a more resistant phenotype, *RR* gene expression was higher in plants treated with KIN 48h or 72hbi than in plants treated with BA. In contrast to KIN, at all hbi tested BA strongly upregulated the expression of *Os C KX 2* which encodes a CK degradation enzyme (Suppl. Fig. S2C).

### Deletion of a gene that encodes a putative CK inactivation enzyme affects endogenous CK levels but not seedling development

To further explore CK roles in rice blast resistance, we characterized resistance and defense responses of mutant plants that have divergent endogenous CK levels (Fig. 2). A putative CK-UDP Glycosyltransferase (*CK-UG T;* LOC_Os02g51930) was identified by previous transcriptomic analyses ^37^. This enzyme family inactivates free base CKs and converts them into storage/transport forms by adding a glucoside group ^38–43^. We obtained homozygous mutants and null-segregant individuals (NS) from two independent insertion lines, *ck-ugt1.1* and *ck-ugt1.2* available in the Dongjin background (Fig. 2A). Whereas no obvious developmental phenotype was associated with the mutation before tillering (<4 week-old plants), a reduced number of tillers was noticed on the adult mutant plants (Fig. 2B; <10 tillers in mutant *vs* >20 in NS). Based on the protein function predicted from the coding sequence mutated, the mutants were hypothesized to be impaired in the CK conversion from the active free-forms to the inactive glucoside forms. Therefore, liquid chromatography-tandem mass spectrometry (LC-MS/MS) was used to quantify CK O- and N-glucosides in 3-week-old healthy plants (stage of experimental inoculation) (Fig. 2C). The level of the *cis* Zeatin-O-glucoside (cZOG), the main form detected, was significantly lower in the *ck-ugt1.1* mutant whereas no differences were found for the other forms (i.e. *cis* Zeatin riboside-O-glucoside (*c*ZROG), Dihydrozeatin-O glucoside (DZOG), *trans* Zeatin-N9-glucoside (*t*Z9G). This analysis supports the hypothesis that the mutants are affected in zeatin-O-glycosylation and that they could be good candidates to further explore the role of endogenous CKs in the rice blast interaction.

**Figure 2.**
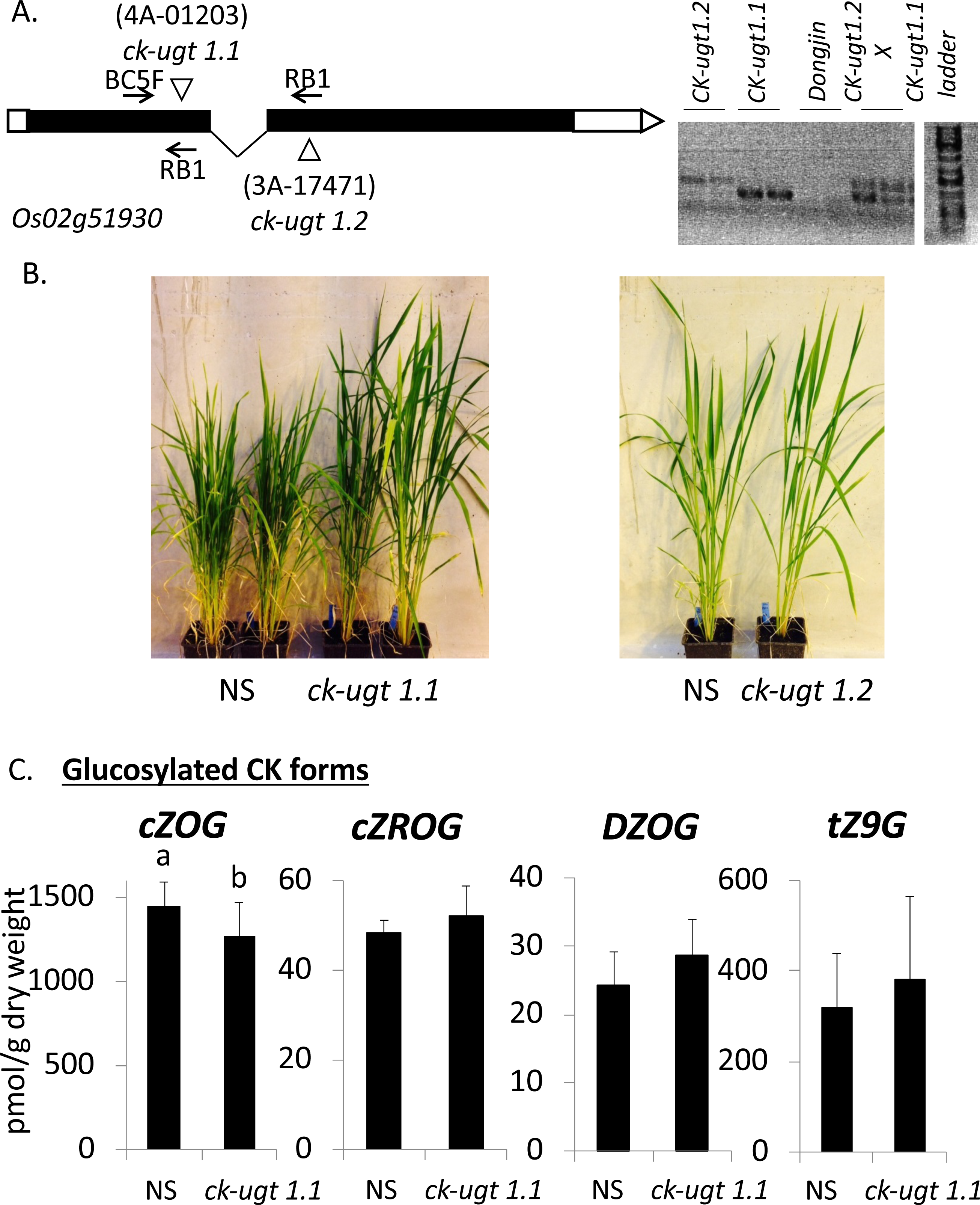
Knock-out mutants, *ck-ugt1.1* show a lower c is ZOG content. (A) Left: Representation of the insertions leading to the disruption of *Os 02g51930*; right: PCR genotyping of the single and double insertion mutant lines (*ck-ugt1.1, ck-ugt 1.2* and *ck-ugt1.1xck-ugt 1.2*). The primers BC55F (on genomic DNA) and RB1 (on T-DNA) were used to genotype mutant plants. (B) The developmental phenotype (reduced tiller number) of *ck-ugt* mutants is visible in adult plants (left; 3-month-old *CK-ugt1.1*) and almost not visible on younger plants (right; 1.5-month-old *CK-ugt1.2*). (C) CK glucoside contents in NS (Null Segregant) plants and *ck-ugt1.1* mutants (t-test; p-value<0.04) measured by LC-MS/MS: cZOG: *cis*-zeatin-O-glucoside; cisZROG: *cis*-zeatin riboside-O-glucoside, DZOG: dihydrozeatin-O-glucoside; tZ9G: *trans*-zeatin-N9-glucoside.

### The KO *ck-ugt* mutation improves blast resistance and enhances plant defenses

To assess the impact of endogenous CKs on blast resistance, we inoculated the *ck-ugt* mutants with the GY11 isolate (Fig. 3 and Suppl. Fig. S5). Based on the number of lesions and on the reduced pathogen growth inside the host tissues, the *ck-ugt1.1* mutants were more resistant compared to the NS control plants (Fig. 3ABC). Similar results were observed with the second independent mutant line (*ck-ugt1.2*), and with the F_2_ plants obtained through the crossing between these two lines, confirming the involvement of the mutation in the resistant phenotype observed (Suppl. Fig. S5). Consistently with the resistance level, the expression of SA-related and the other defense-marker genes was higher in the healthy and infected *ck-ugt1.1* mutants compared to the NS individuals (Fig. 3D, Suppl. Fig. S6). Indeed, excepted *Os WR K Y 45*, this higher expression of defense markers was no longer visible 48h after inoculation. These results suggest that modulating endogenous CK homeostasis enhances rice resistance likely by inducing SA-dependent defenses even in absence of pathogens.

**Figure 3.**
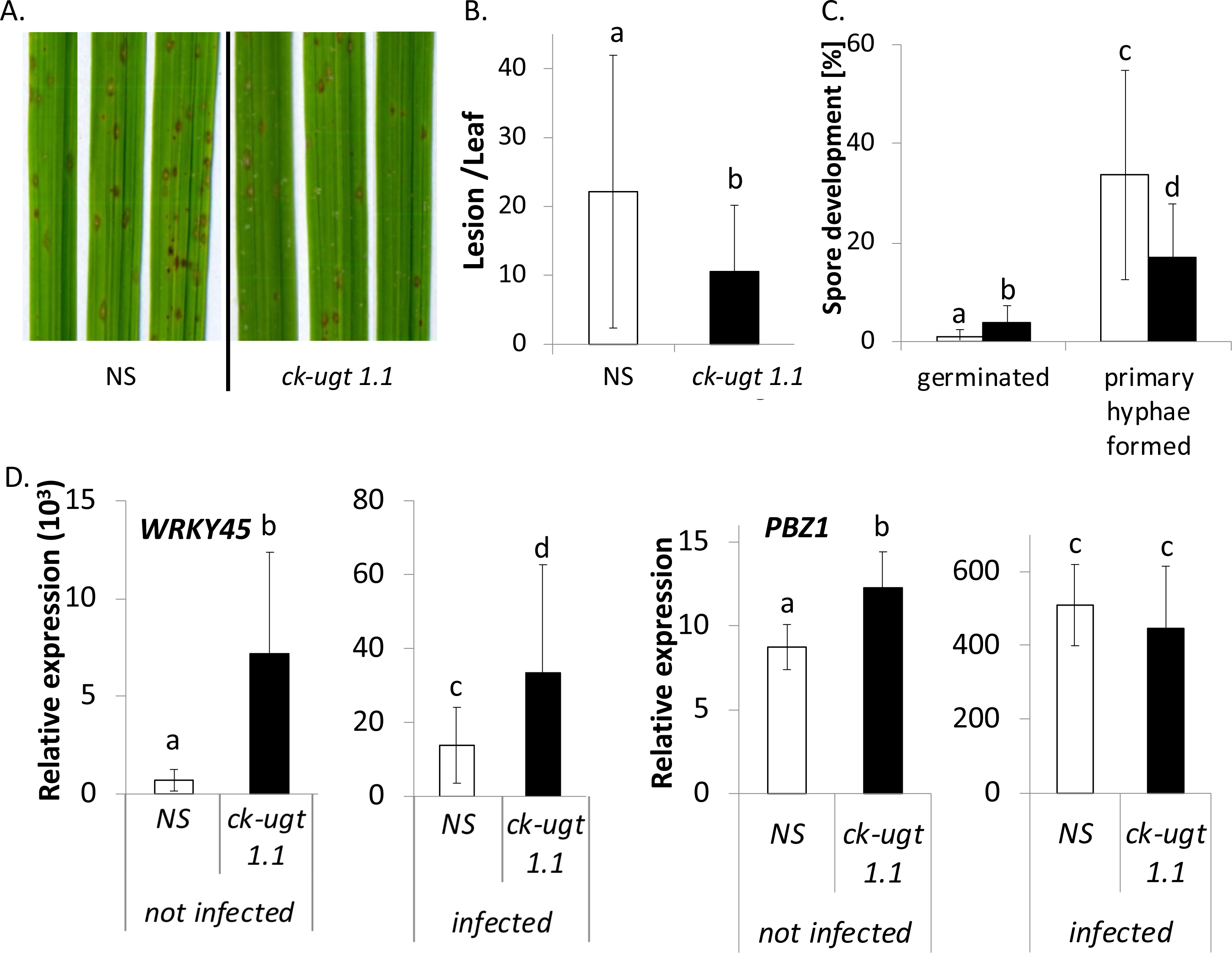
ck-ugt1.1 mutant plants are more resistant to M. oryzae. (A) Symptoms caused by the *M. oryzae* strain GY11 on NS (null segregant) plants and *ck-ugt1.1* mutants observed 6 days after inoculation. (B) Symptom quantification. The values represent the mean and SD from five to eight biological replicates each composed of 10 plants (t-test, p-value < 0.04). (C) Microscopic observations of spore germination and penetration 72h post inoculation in NS plants (white bars) and *ck-ugt1.1* mutants (black bars). The data presented is the mean and SD of three biological replicates (>50 infection sites/replicate, t-test, p-value < 0.03). (D) Relative expression of defense-marker genes, *Os WR K Y 45* and *PBZ 1*, in not infected and infected (72h post inoculation) NS (white bars) and *ck-ugt1.1* mutant plants (black bars). The values presented are the means and SD calculated from four to six independent replicates composed of 10 plants (t-test, p-value < 0.03). The transcriptional regulation was evaluated by quantitative RT-PCR using the *Actin* gene for normalization.

## Discussion

Applying exogenous CKs mimicking compounds is a standard approach to investigate their roles in plant immunity ^6,9,22,44–46^. By this approach Jiang et al, (2013) showed *in vitro* that CKs and SA act synergistically to induce defense marker genes but no resistance phenotype was obtained *in planta* ^9^. Likewise, we also did not observe any difference in blast resistance with a 50µM KIN treatment 24hbi (Suppl. Fig. S2AB). Nonetheless we demonstrated that CKs contribute to rice immunity in a compound-, dose- and time-dependent manner (Fig. 1; Suppl. Fig. S1, S2, S3). This is similar to the results of Argueso et al., (2012) who showed that *Arabidopsis thaliana* resistance to *Hyaloperonospora arabidopsidis* varies according to CK doses applied 2 days before inoculation. Thus we established that 50µM KIN 48hbi were the conditions leading to the most reproducible rice blast resistance phenotype and confirmed *in planta* that SA and CKs synergistically enhance defense genes (Fig 1D, Suppl. Fig. S1D, S4). Also, the variability of the CK-induced resistance depending on compounds, doses and timing of application coincides with differences in CK-marker genes expression (Suppl. Fig. S2C). In agreement with the literature ^47^, this suggests that the molecule and dose could lead to different CK responses with different effects on the resistance levels (Suppl. Fig. S2). Indeed, KIN or BA have varying CKs activity already reported in other cellular processes ^48^. Altogether, exogenous application of plant hormone-mimicking compounds may lead to artefactual effects and misinterpretations but remain a sound tool when mutating endogenous hormonal pathways causes severe developmental defects ^8–10^. Here, to further evaluate the role of endogenous CKs in rice resistance, we characterized rice *ck-ugt* insertion mutants in the gene LOC_Os02g51930. This gene encodes a putative CK-UDP Glycosyltransferase (*CK-UG T;* LOC_Os02g51930) and is highly expressed during infection with the virulent isolate FR13 ^37^. The LC-MS/MS CK quantification in the mutants revealed a ∼15% decrease of *c*ZOG levels, the most abundant glucosylated CK forms (Fig. 2C). This phenotype is similar to the *Os ZOG 1* rice mutants characterized by Shang et al., (2016) showing ∼17% reduction in *c*ZOG levels ^41^. Even though this was a slight difference, the authors noticed it sufficient to have physiological impacts in rice ^41^. Moreover, both genes, *CK-UG T* and *Os ZOG 1* (LOC_Os04g20330 ^41^), share 25% identity supporting the idea that *CK*-*UG T* also encodes a zeatin-O-glycosyltransferase. In the literature, the CK glucosides are described as inactive storage/transport forms that are converted back into the active free bases or ribosides ^43,49^. Since the *ck-ugt* mutants have altered endogenous CK profiles suggesting an impaired CK inactivation pathway but do not present obvious developmental defects at the stage of infection, they were good candidates to confirm the involvement of rice CKs in immunity (Fig 2B). The *ck-ugt* mutation indeed conferred a higher level of resistance with a reduced pathogen growth inside host tissues (Fig 3AB). These results correlate with a higher constitutive defense gene expression in mutant plants compared to the control ones prior to inoculation (Fig. 3D and Suppl. Fig. S6). Depending on the defense markers, the difference in expression level was not always persistent 48 hours after inoculation (Fig 2). This suggests an efficient priming effect happening in the *ck-ugt* mutants that compromises pathogen invasion at the early stages (Fig. 3D and Suppl. Fig. S6). However, the adult *ck-ugt* mutants showed a reduced tillering that may be due to a cost of expressing defenses in the absence of a pathogen threat (Fig 3D). Moreover, CKs are involved in many plant physiological processes and it cannot be excluded that the *ck-ugt* mutation have deleterious effects under other abiotic or biotic stresses involving pathogens with different trophic behaviours ^5,50,51^. Also, CKs may indirectly promote rice resistance by impacting *M. oryzae,* which is known to require CKs for its full virulence ^9,24^. Indeed, several recent studies on tomato and *Botrytis cinerea* revealed interesting roles of CKs in plant-pathogenic filamentous fungi interactions ^52,53^. Gupta et al., (2021a) showed that CKs impact fungal cytoskeleton structure and cellular trafficking ^54^. This warrants additional microscopic analyses of *M. oryzae* spore development upon infection to better understand what is happening on pathogen’s physiology. Additionally, CKs are emerging as key interkingdom communication factors affecting host microbiome structure with impact on resistance ^55^. This widens the range of future disease management strategies, especially for staple crops such as rice in which CKs roles on immunity remain mostly elusive.

## Materials and Methods

### Plant Growth

Before CKs treatment and/or inoculation, rice (*O ryz a sativa*) plants were grown for three weeks as described previously by Faivre-Rampant *et al.* (2008) ^56^.

### Fungal Growth and Inoculation

Two *Magnaporthe oryzae* isolates were used, GY11 and FR13. Both were applied to evaluate Nipponbare resistance after exogenous CK treatments whereas only GY11 was virulent on the Dongjin background. Fungal isolates were grown on rice flour agar for 10 days ^57^ then a suspension of fungal conidiospores (5×10^4^ sp/mL in 0.5% gelatin) was sprayed on 3-week-old rice plants. To evaluate the resistance level, symptoms were quantified on 6-10 leaves from 3-4 independent biological replicates (each containing 10 plants). All experiments were at least repeated 3 times.

### Exogenous CKs Treatment

The exogenous CKs assays were performed on the rice cultivar Nipponbare. Kinetin (KIN; 10, 50, 100 and 500µM) and Benzyladenine (BA; 50µM) were dissolved in 50% ethanol to obtain solutions diluted at least x1000 to be sprayed on plants 24h, 48h and 72h prior to fungal inoculation. The equivalent of 50% ethanol was diluted into water to be used as control (mock) treatments. Since CKs interact with plant nitrogen metabolism and that nitrogen supply impacts rice blast resistance ^58^, we established a standard protocol to reproduce their effects on rice immunity: after 3 weeks of growth, day 1 plant fertilization, day 3 CKs sprays, day 5 inoculation with fungal spores.

### Endogenous CKs mutants and endogenous CKs levels measurements

Two *CK-UG T* knock-out mutant lines in the Dongjin background, *ck-ugt1.1* and *ck-ugt1.2 (Knock-Out - KO- and* Null Segregants - NS) and the F2 *ck-ugt1.1*×*ck-ugt1.2* individuals were used in the experiments. The *ck-ugt* mutants were obtained through T-DNA insertion leading to the disruption of *CK-UG T* gene (MSU accession: *Os 02g51930*, alias RAP accession *Os 02g0755900*). The BC55F (on genomic DNA) and RB1 (on T-DNA) primers were used to genotype mutant plants (Fig. 2.).

The levels of CK glucosides (O- and N-glucoside derivatives of tZ (tZOG, tZ9G, tZROG), cZ (cZOG, cZ9G, cZROG) and DZ (DZOG, DZ9G and DZROG)) were measured using liquid chromatography-tandem mass spectrometry (LC-MS/MS) in 3-week-old healthy control plants (NS) and *ck-ugt* KO rice mutants as previously described ^18,24^.

### Confocal Observations of Fungal Penetration

To evaluate cellular fungal penetration, leaves were harvested upon infection, fixed and stained as described by Ballini et al., (2013) ^58^. Three leaves per time point and condition were collected from 4 independent biological replicates (each containing 10 plants). Each experiment was at least repeated twice.

### RNA Extraction and Quantitative RT–PCR Analysis

For gene expression analysis, an inoculum of 2×10^5^ sp/mL in 0.5% gelatin was used. RNA extraction from inoculated and control rice plants was performed 8h, 24h, 48h, 72h, and 96h post infection as previously described ^59^. Quantitative PCR was performed using LC 480 SYBR Green I Master Mix (Roche, Basel, Switzerland) and a LightCycler 480 Real-Time PCR instrument (Roche). cDNA amplification was performed as follows: 95°C for 10 min; 40 cycles of 95°C for 15s, 60°C for 20s and 72°C for 30s; then 95°C for 5 min and 40°C for 30s. The expression of SA-related defense-marker genes, *WR K Y 45* and *PBZ 1*, five other defense-marker genes (*POX 223*, *PR 10*, *PR 5*, *SP L 7* and *CE B iP*), CK degradation gene (*Os C KX 2*) and three *Os R R* genes (*Os R R 2*, *Os R R 3*, *Os R R 6*) were analyzed using forward and reverse primers indicated in Table S1. The *Actin* gene was used for data normalization. For each time point, leaves were collected from 4 independent biological replications (each containing 10 plants) and each experiment was at least repeated twice.

### Data and Statistical Analysis

Data analyses and representations were performed with Microsoft Excel© and the R packages tidyverse ^60^, tidyr ^61^, dplyr ^62^ and ggplot2 ^63^. The effect of plant genotype (NS and *ck*-*ugt* mutants) and CK pretreatment (compound, dose and timing of application) on the number of lesions, spore germination and hyphal growth, was determined using t-tests at the p value ≤ 0.05 (R, package easystats ^64^). The relative expression of CK metabolism- and defense-related marker genes was respectively determined by using t-tests and one-way analysis of variance followed by a Tukey’s Post hoc test, p value ≤ 0.05 (R, package easystats ^64^).

## Acknowledgements

We thank Bastien Cayrol for multiplying the *ck-ugt* mutants and for designing the genotyping primers.

## Author Contributions

Conceived and designed the experiments: JBM EC. Performed the experiments: EC AK SB CRM JBM. Analyzed the data: EC AK JBM. Contributed reagents/materials/ analysis tools: EC AK JBM. Wrote the paper: EC JBM AK NRJE.

## ABBREVIATION

BA: BenzylAdenine
CEBiP: Chitin Elicitor Binding Protein
CK: Cytokinins
CKX: Cytokinin oxidase
cZOG: *cis*-Zeatin-O-glucoside
cZROG: *cis*-Zeatin riboside-O-glucoside
DZOG: Dihydrozeatin-O-glucoside
Hai: hours after inoculation
Hbi: hours before inoculation
KIN: Kinetin
LC-MS/MS: Liquid Chromatography tandem Mass-Spectrometry
NS: Null Segregant
Os: *Oryza sativa*
PBZ1: Probenazol 1
POX223: Class III plant Peroxidase 223
PR5: Pathogenesis-Related 5
RR: Response Regulator
SA: Salicylic Acid
SPL7: Spontaneous Lesion 7
tZ9G: *trans*-Zeatin-N9-glucoside
UDP: Uridine diphosphate
UGT: UDP-GlycosylTransferase
WRKY45: WRKY domain containing transcription factor 45

