## Supplementary material for "Cytokinin-induced immunity enhances rice blast resistance": suppl. data

Figure S1.

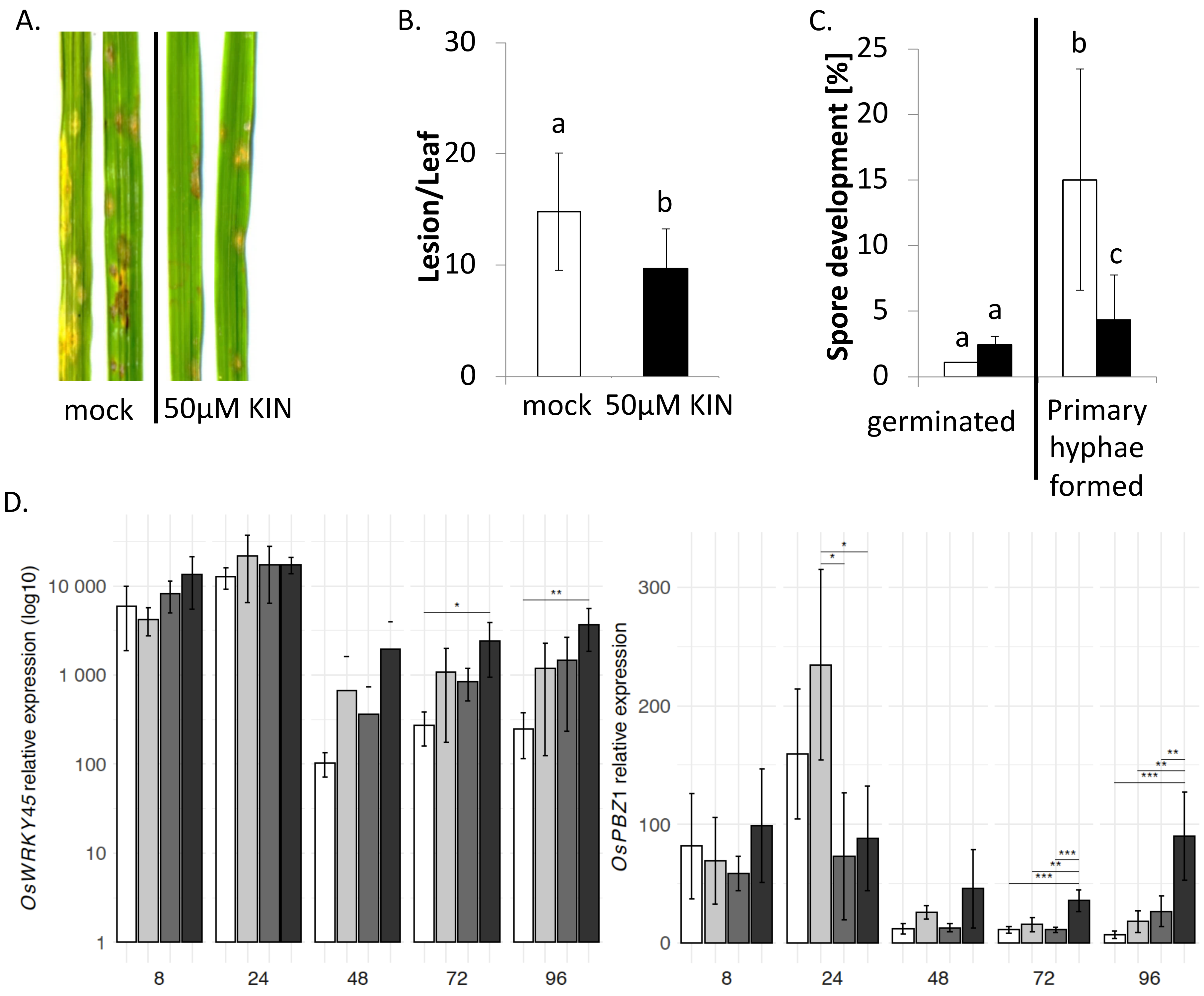
A.
mock
50µM KIN
B.
Lesion/Leaf
0
10
20
30
a
b
mock
50µM KIN
C.
Spore development [%]
0
5
10
15
20
25
a
a
b
c
germinated
Primary hyphae formed
D.
OsWRKY45 relative expression (log10)
1
10
100
1 000
10 000
8
24
48
72
96
OsPBZ1 relative expression
0
100
200
300
8
24
48
72
96

**Figure S1. Kinetin enhances resistance against the FR13 *M. oryzae* isolate.**

(A) Symptoms caused by the *M. oryzae* strain FR13 on Nipponbare plants treated with 50µM of KIN 48hbi and on the controls (mock) observed 5 days after inoculation. (B) Symptom quantification. The values represent the mean and SD from six biological replicates each composed of 10 plants (t-test, p-value = 0.02). (C) Microscopic observations of spore germination and penetration 96h post inoculation in plants pretreated with 50µM of KIN (black bars) 48hbi or with the mock solution (white bars). The data presented is the mean and SD of three biological replicates (with 83 infection sites/replicate in average, t-test, p-value = 0.02). (D) Relative expression of the defense-marker genes, *OsWRKY45* and *PBZ1*, in mock-treated plants sprayed with 0.5% gelatin (white bars), in KIN-treated plants sprayed with 0.5% gelatin (light grey bars), in mock-treated infected plants (dark grey bars) and in KIN-treated infected plants (black bars) at 8, 24, 48, 72 and 96h after inoculation (hai). The transcriptional regulation was evaluated by quantitative RT-PCR using the Actin gene for normalization. The values presented are the means and SD calculated from four independent replicates composed of 6 plants. Asterisks indicate significance in a one-way analysis of variance with a Tukey post hoc test, $p < 0.04$). For all the experiments, KIN 50mM in 50% EtOH diluted x1000 (KIN 50µM final) or the equivalent 50% EtOH control mock solution were applied to plants 48hbi. Inoculated plants were sprayed with spore suspension (in gelatin 0.5%) and uninfected controls with gelatin (0.5%) only.

Figure S2.

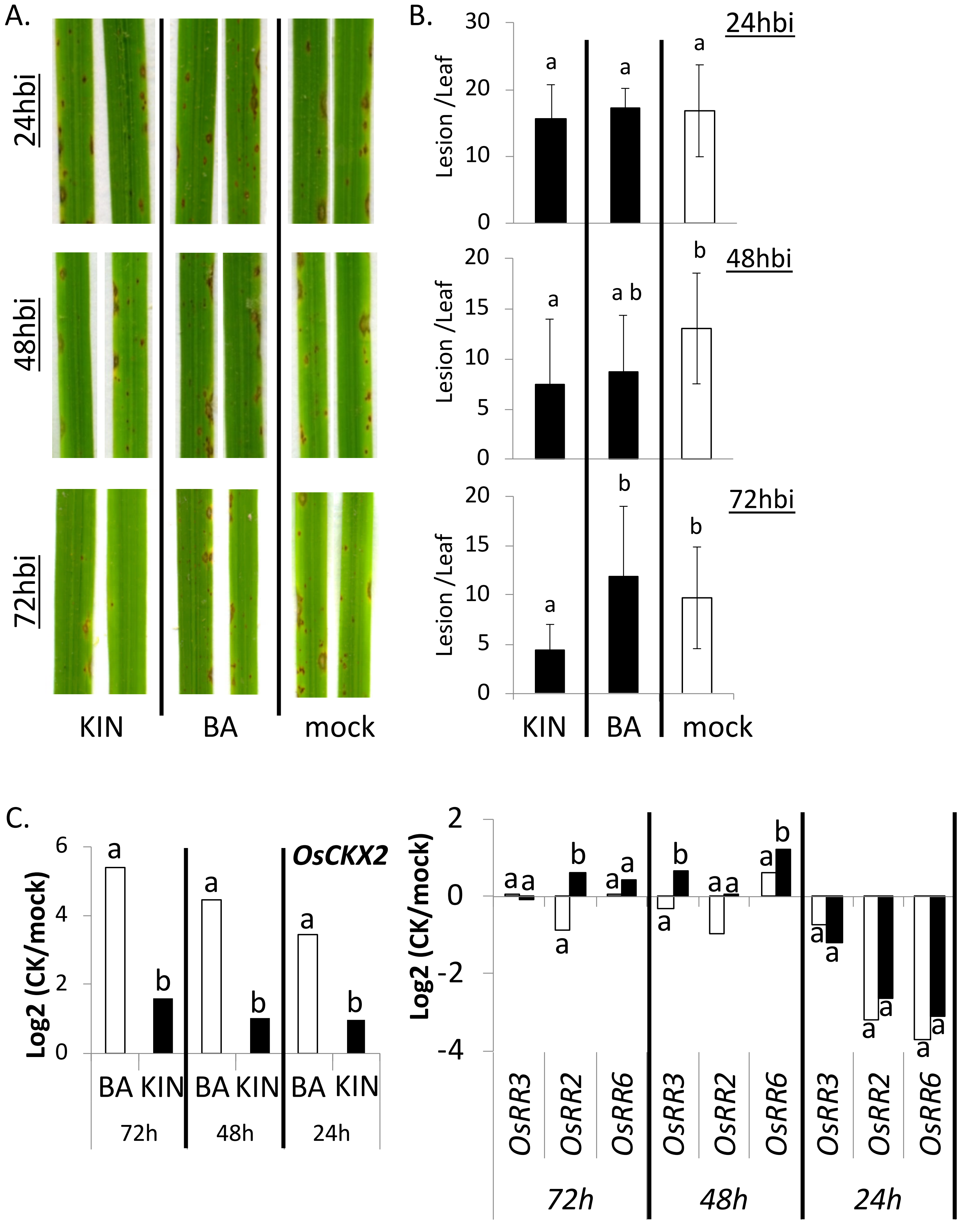
A.
24hbi
48hbi
72hbi
KIN
BA
mock
B.
24hbi
48hbi
72hbi
Lesion /Leaf
KIN
BA
mock
C.
OsCKX2
Log2 (CK/mock)
BA KIN BA KIN BA KIN
72h
48h
24h
OsRR3
OsRR2
OsRR6
72h
48h
24h

**Figure S2. Kinetin and Benzyladenine affect rice blast resistance and marker gene expression differently.**

(A) Symptoms caused by the *M. oryzae* strain GUY11 on Nipponbare plants treated with 50µM of KIN or Benzyladenine (BA), 24h, 48h, and 72hbi and in control (mock) plants observed 6 days after inoculation. (B) Symptom quantification. The values represent the mean and SD from at least six biological replicates each composed of 10 plants (t-test, p-value < 0.02). (C) Relative gene expression 48h post inoculation in plants pre-treated with 50µM BA (white bars) or KIN (black bars) 72h, 48h or 24hbi (t-test, p-value < 0.04). The transcriptional regulation of the CK metabolism-related genes: *OsCKX2* (left) and three *OsRRs* (right) were evaluated by quantitative RT-PCR using the *Actin* gene for normalization. The values presented are the Log2 ratios of the means (CK/mock) of four independent replicates composed of 6 plants.

Figure S3.

A.

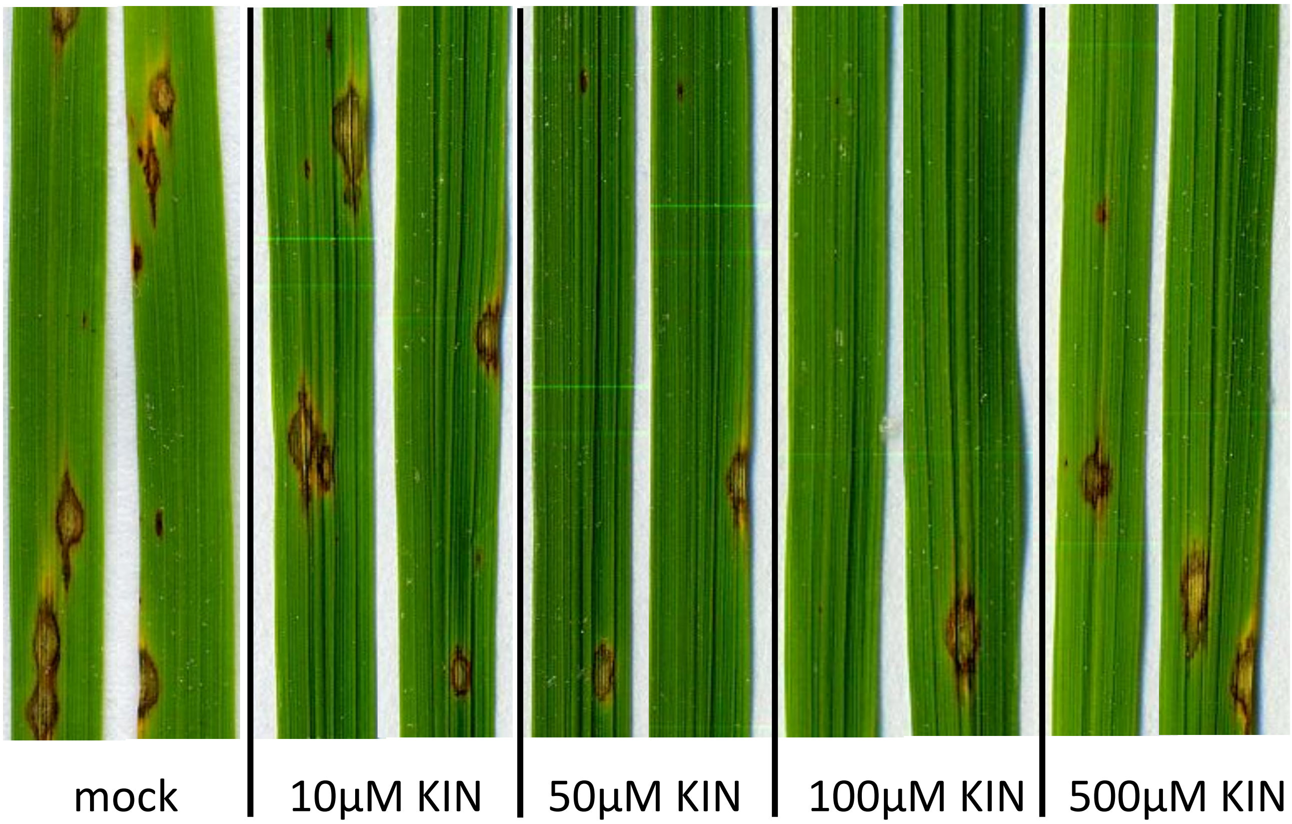
mock
10μM KIN
50μM KIN
100μM KIN
500μM KIN

B.

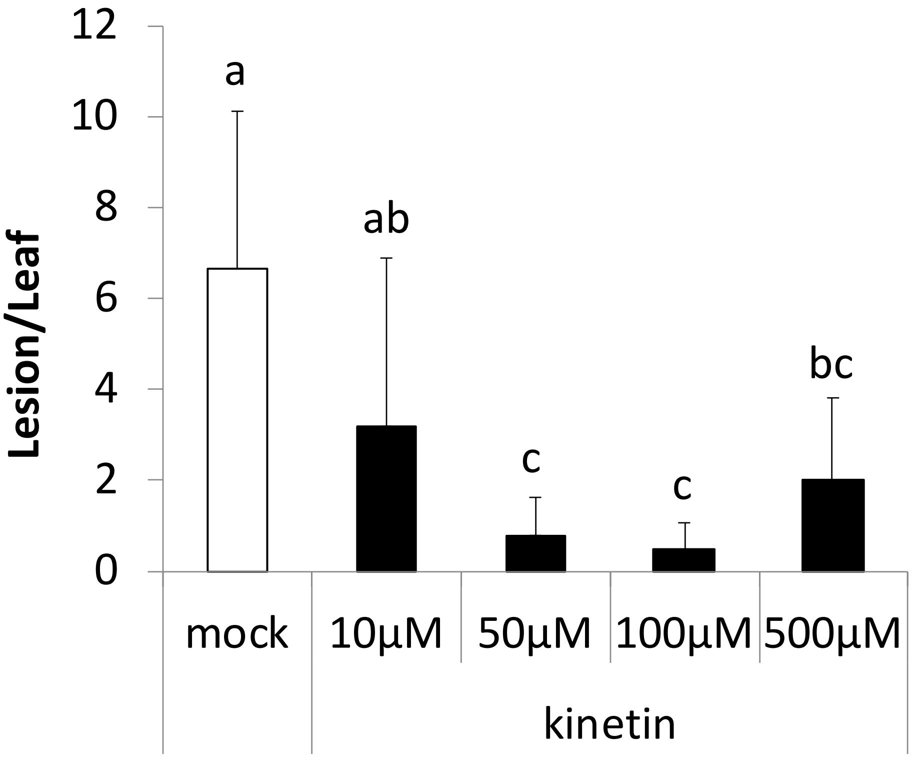
12
10
8
6
4
2
0
Lesion/Leaf
a
ab
c
c
bc
mock
10μM
50μM
100μM
500μM
kinetin

**Figure S3. Kinetin induces rice blast resistance in a dose-dependent, biphasic manner.**

(A) Symptoms caused by the *M. oryzae* strain GY11 on Nipponbare plants treated with different concentrations of KIN 48hbi and on plants treated with the mock solution observed 6 days after inoculation. (B) Symptom quantification. The values represent the mean and SD from at least six biological replicates each composed of 10 plants (t-test, p-value = 0.03), in mock-treated plants (white bars) and in KIN-treated plants (black bars).

Figure S4.

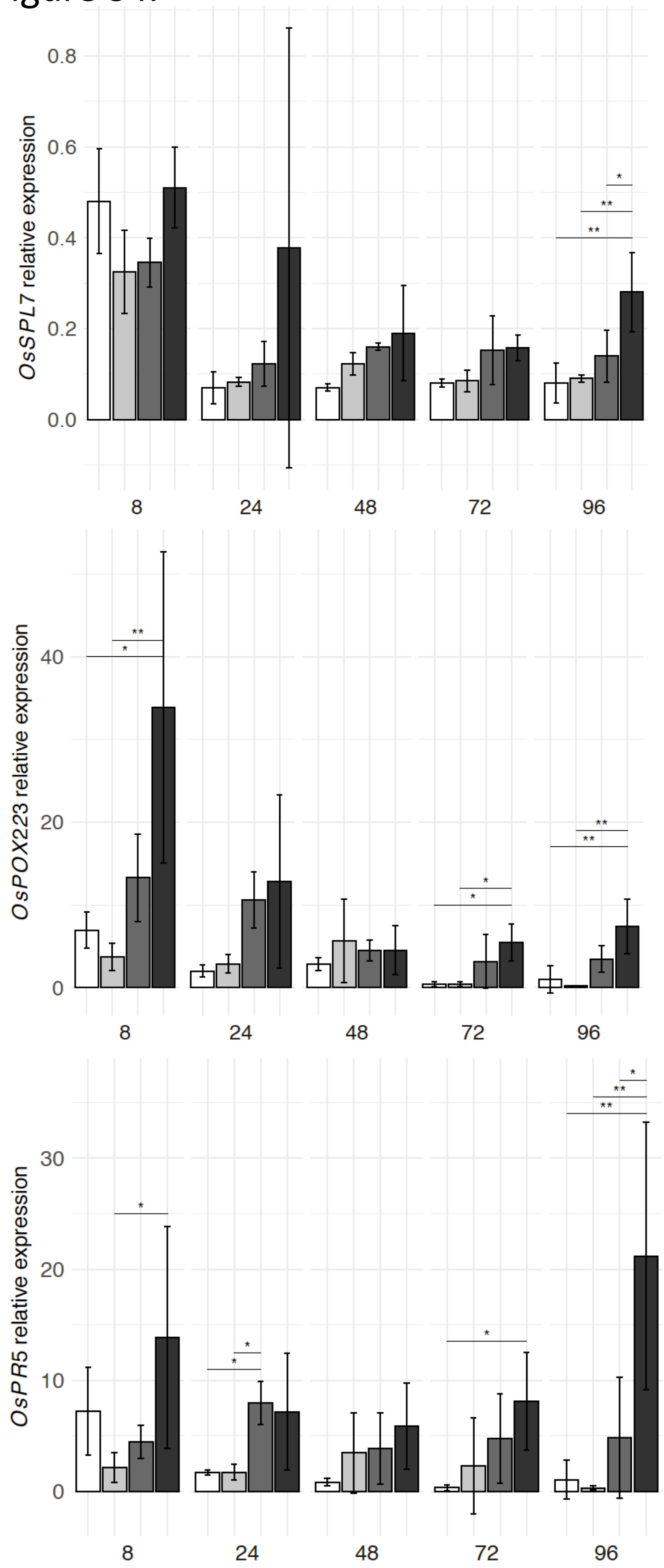

OsSPL7 relative expression
0.8
0.6
0.4
0.2
0.0
8
24
48
72
96
OsPOX223 relative expression
40
20
0
8
24
48
72
96
OsPR5 relative expression
30
20
10
0
8
24
48
72
96

**Figure S4. Relative expression of defense marker genes upon infection is higher in KIN-treated plants.**

Relative expression of the defense-marker genes in mock-treated plants sprayed with 0.5% gelatin (white bars), in KIN-treated plants sprayed with 0.5% gelatin (light grey bars), in mock-treated infected plants (dark grey bars) and in KIN-treated infected plants (black bars) at 8, 24, 48, 72 and 96h after inoculation (hai). The transcriptional regulation of defense marker genes, *SPL7*, *POX223* and *PR5*, was evaluated by quantitative RT-PCR using the *Actin* gene for normalization. The values presented are the means and SD calculated from four independent replicates composed of 6 plants. Asterisks indicate significance in a one-way analysis of variance with a Tukey post hoc test, $p < 0.04$). For all the experiments, KIN 50mM in 50% EtOH diluted x1000 (KIN 50μM final) or the equivalent 50% EtOH control mock solution were applied to plants 48h before inoculation. Inoculated plants were sprayed with spore suspension (in gelatin 0.5%) and uninfected controls with gelatin (0.5%) only.

Figure S5

A.

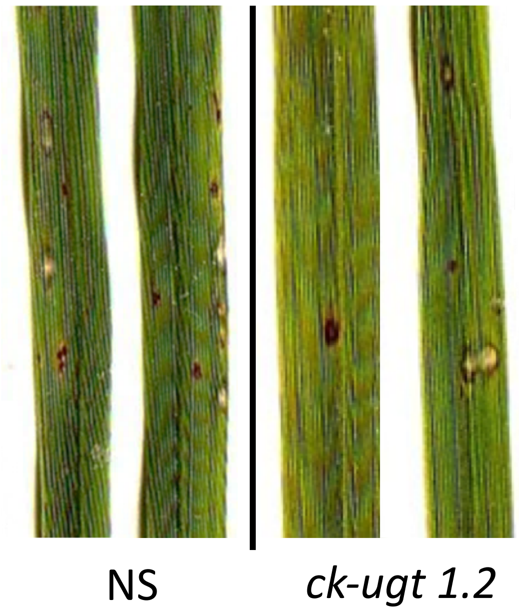


B.

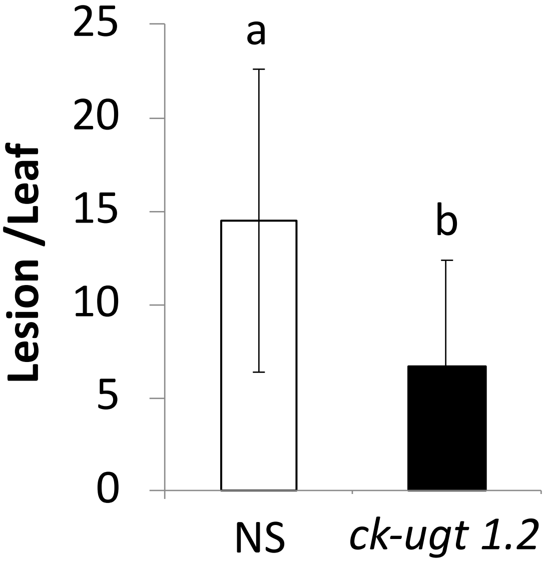


C.

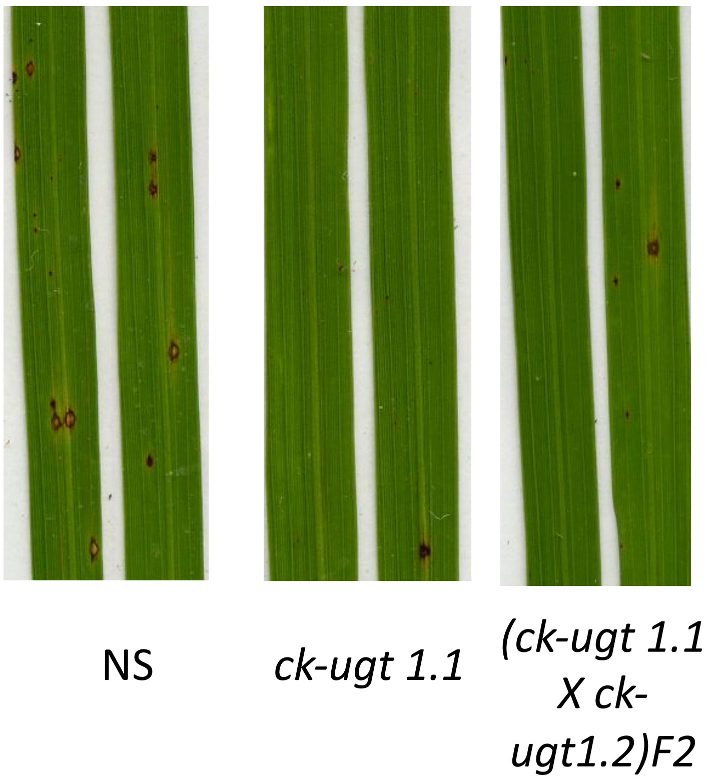

**Figure S5. *ck-ugt1.2* and (*ck-ugt1.1 x ck-ugt1.2*)$F_2$ mutant plants are more resistant to *M. oryzae*.**

(A) Symptoms caused by the *M. oryaze* strain GY11 on *ck-ugt1.2* mutants and null segregant (NS) control plants observed 7 days after inoculation. (B) Symptom quantification. The values represent the mean and SD from six biological replicates each composed of 10 plants (t-test, p-value < 0.04). (C) Symptoms caused by the *M. oryaze* strain GY11 on null segregant (NS) control plants, on *ck-ugt1.1* mutants (*ck-ugt1.1*x*ck-ugt1.2)*$F_2$ observed 7 days after inoculation. For each time point, the mean values and SD of at least 4 biological replicates composed of at least 10 plants are presented.

Figure S6.

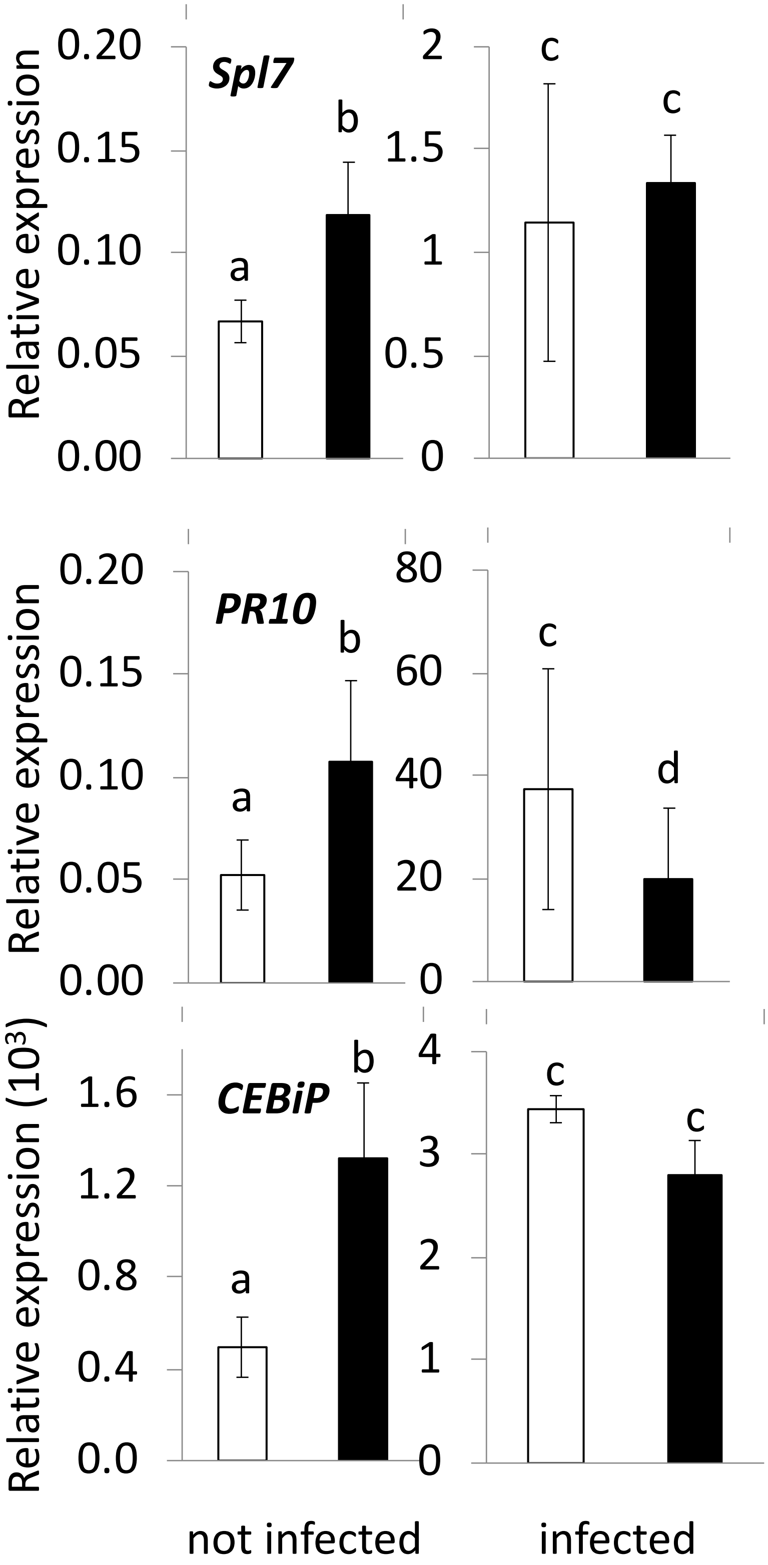

Spl7
PR10
CEBiP
Relative expression
Relative expression
Relative expression (10$^3$)
0.20
0.15
0.10
0.05
0.00
2
1.5
1
0.5
0
80
60
40
20
0
1.6
1.2
0.8
0.4
0.0
4
3
2
1
a
b
c
d
not infected
infected

**Figure S6. Defense marker gene expression is higher in *ck-ugt* mutants in absence of the pathogen.**

The transcriptional regulation of defense marker genes, *Spl7*, *PR10* and *CEBiP*, was evaluated by quantitative RT-PCR using the *Actin* gene for normalization in *ck-ugt1.1* (black bars) and in null segregant plants (white bars). The values presented are the means and SD calculated from four independent replicates composed of 3 plants (t-test, p-value <0.05).

### Figure S7

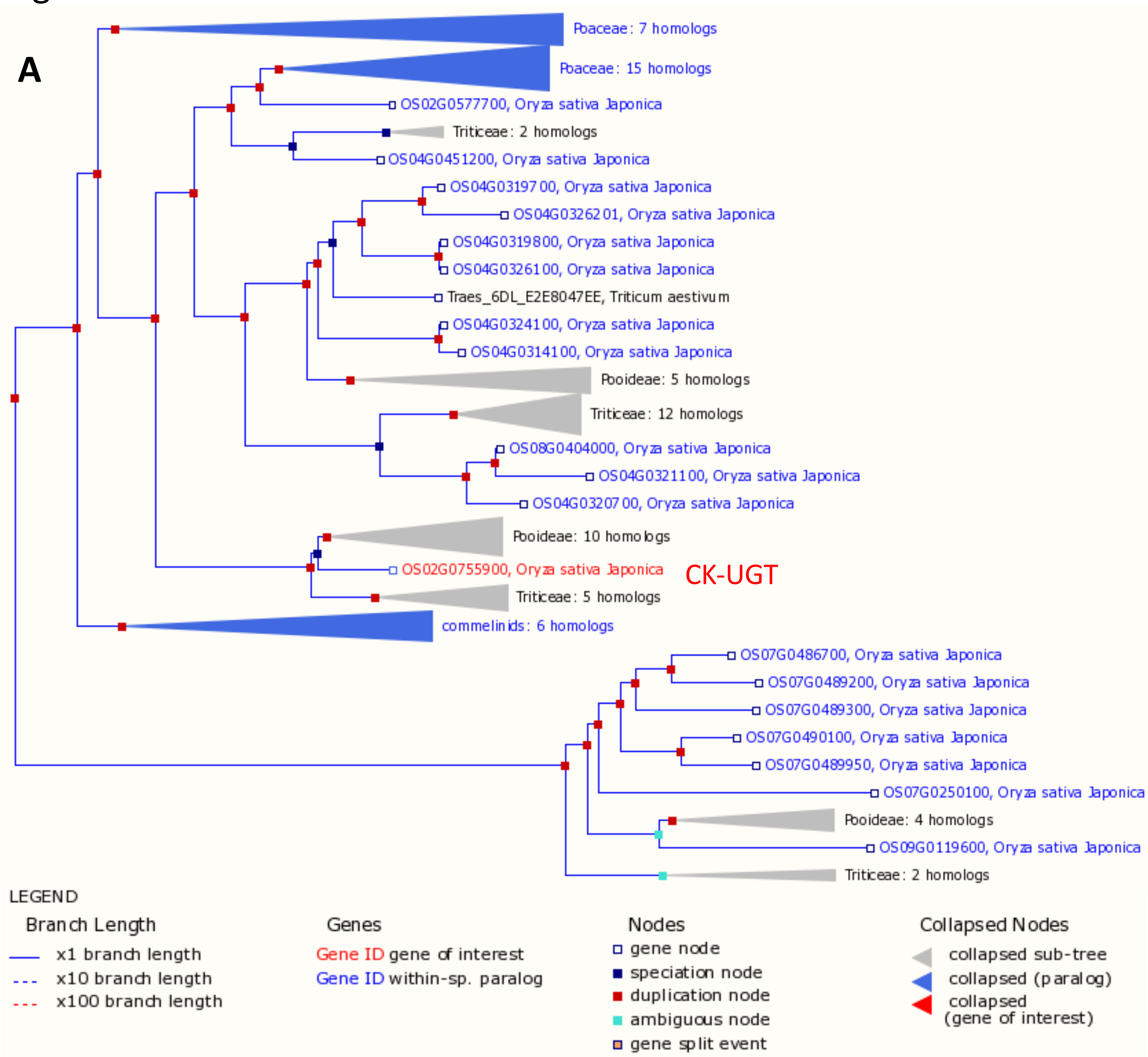


**B**

```
               1       10        20        30        40        50        60        70        80        90        100       110       120       130
               |--------+---------+---------+---------+---------+---------+---------+---------+---------+---------+---------+---------+---------|
Os04g20330_OsZOG1     MPSDGSFRIVLFPFPAQGHFSAFLSLAAHLHDAQPTADITIVSTPRNVEDLRRRSSSQTRYLRFHALPFAPAEHGLPGDIESTDAVPLLHFITLFEATESRSLQDSFDSFVRDLITDAGADGARVCV
Os02g51930_CK-UGT  MGSLGAAGKPHAVCMPYPAQGHITPMLNVAKLLHARG--FDVTFVNTEYNHARLVR-TRGEAAVAGAPGFRFATIPDGLP---PSDDDDVTQDIPSLCRSTKETCLAP-FRRLLAQLNDPATGHPPVTCV
        Consensus  ...$gaaGkfraVcmP%PAQGHisamLnlAahLHaaq..aD!TiVnTerNharLrR.srg#aayarahalrFApaedGLP...eSdDadplqdiisLcraTeercLad.Frrlla#LnddAgadgartCV

               131  140       150       160       170       180       190       200       210       220       230       240       250       260
               |--------+---------+---------+---------+---------+---------+---------+---------+---------+---------+---------+---------+---------|
Os04g20330_OsZOG1  IADPFLAWTTDVARRRGAAHAIFVSCGAFGSVVFHSLWNHLPHLRAPGDDAFCLP---------DHPEVTVHRSQLPPYLLHADGTDRWSAHHR-RQTSAGYDTDAILISTMEELETTGLRMLRKTMGVP
Os02g51930_CK-UGT  VSDVVMGFSMAAAKELGLPYVQLWTASSISYLGYRHYRLLMERGLAPLKDVDQLTNGYLDTPVEDVPGLRNMRIKDFPSFIHTTNPEEYMVGYVIEETERCKDASAIIVNSFGDLEGEAVAAMEALGLPK
        Consensus  !aDpf$afsmaaArrrGaahaqlwsagaigslg%rhlrnh$ergrAPgdDadcLp.........DhPelrnhRiqdfPsliHadnp#rwmaghr.r#TeackDadAIi!nsme#LEgealaa$ralggpk

               261  270       280       290       300       310       320       330       340       350       360       370       380       390
               |--------+---------+---------+---------+---------+---------+---------+---------+---------+---------+---------+---------+---------|
Os04g20330_OsZOG1  VYPIGPL--VRRRTEHSDHIGD------HNDDDVKRWLDTREERSVLYISFGSNNSLRPDQMVDLAMALELTGRPFIWAIRPPFGFDIETTNGREFSAEWLPEGFEERMRAKNIGLLIHGWAPQVSILAH
Os02g51930_CK-UGT  VYTLGPLPLVARKDPPSPRRSSIRLSLWKEQEECLQWLDGKEAGSVVYVNFGSITVMTNEQLVEFAWGLANSGREFLWIVRRDLVKGDTAVLPPEFLAETAERG-----------LMASWCPQQDVLNH
        Consensus  VYpiGPL..VaRrdehSdrrgd......h####clrWLDgrEarSVlY!nFGSnns$rn#Q$V#lAmaLansGReFiWa!Rrdlgkddeatngr EFlAEtaerG...........LiagWaPQqd!LaH

               391  400       410       420       430       440       450       460       470       480       490       500       510  515
               |--------+---------+---------+---------+---------+---------+---------+---------+---------+---------+---------+---------+----|
Os04g20330_OsZOG1  ASTGAFLSHCGWNSVLESMAHGVPIIAWPLTADQFFNAQ-MLEEWGACVEVSRGNWPDSPALERERVVEVVEMVMGITAKADKIRQSVKQIQGMIGRTLEDGGSSKTALEEFLKLHGHIMLMKC
Os02g51930_CK-UGT  PAVGAFLTHSGWNSTLESLAAGVPVISWPFFADQQTNCRYQCNEWGVGMEI------DSNVKRGAVACLIAELMEG--QKGKEMRRKAEEWREKAIRAAKPGGSSHRNFEELVR---HVLLAKES
        Consensus  aatGAFLsHcGWNStLES$AaGVP!IaWPlfADQqfNar.qc#EWGacmE!......DSnalrrarace!aE$meG..aKadeiRrkae#irekagRaaedGGSShralEEllr...H!$LaKc.
```

**Figure S7. *CK-UGT* and *OsZOG1* genes are distantly related.**

(A) The *CK-UGT* gene (MSU accession: Os02g51930, alias RAP accession Os02g0755900) is quite unique among rice homologous genes and has homologs in other monocots. Screenshot from Gramene website. (B) The CK-UGT and OsZOG1 proteins share 27.68% identity (conserved residues in red).

### Table S1.

| Accession | Annotation | Sequence |
| --- | --- | --- |
| Os12g36880 | *PBZ1* | F-AGGCATCAGTGGTCAGTAGAG |
| | | R-CGGGTCTTGTATGTGCTTCC |
| Os05g25770 | *OsWRKY45-1* | F-ACGACGAGGTTGTCTTCGATCTG |
| | | R-GCCCGTGTCCATCCATGATTCTTC |
| Os07g48020 | *POX223* | F-CTCAGCTGCTCCAAGGTGAA |
| | | R-TGTGGCCCGTTTATGTCGTC |
| Os12g36850 | *PR10* | F-CAGATGATCGAGGCGTACCT |
| | | R-CCACGCCACAGTAACATGAC |
| Os12g43430 | *PR5* | F-AGCCAGGACTTCTACGACCT |
| | | R-GCGTGTGTCTTGGTGTTGTC |
| Os05g45410 | *SPL7* | F-CGGATTAGAGGCTTGCGTGTTAC |
| | | R-GCACAGTAGTCAGCGGATAGAAC |
| Os03g04110 | *CEBiP* | F-CACTTGTACGGCTGCTTGAA |
| | | R-GGAAGGTGGGAAGTCCATTC |

**Table S1. Primers of rice defense marker, CKX and RR genes used for qRT-PCR.**
